# GPER-dependent estrogen signaling increases cardiac GCN5L1 expression and MCAD activity

**DOI:** 10.1101/2021.09.20.461099

**Authors:** Janet R. Manning, Dharendra Thapa, Manling Zhang, Michael W. Stoner, John C. Sembrat, Mauricio Rojas, Iain Scott

## Abstract

Reversible lysine acetylation regulates the activity of cardiac metabolic enzymes, including those controlling fuel substrate metabolism. Mitochondrial-targeted GCN5L1 and SIRT3 have been shown to regulate the acetylation status of mitochondrial enzymes, which results in alterations to the relative oxidation rates of fatty acids, glucose, and other fuels for contractile activity. However, the role that lysine acetylation plays in driving metabolic differences between male and female hearts is not currently known. In this study, we report that estrogens induce the expression of GCN5L1 via GPER agonism in cardiac cells, which increases the enzymatic activity and acetylation status of the fatty acid oxidation enzyme medium chain acyl-CoA dehydrogenase (MCAD).

## INTRODUCTION

Improved understanding of the physiological and metabolic differences between men and women may allow us to develop new therapies that can address sex-based disparities in cardiac disease treatment outcomes. Sex hormones testosterone and estrogen, as well as chromosomal effects, may contribute to sex-based differences. Pre-menopausal women exhibit increased estrogen levels relative to men and post-menopausal women, which results in greater activation of estrogen receptors in the myocardium. These are comprised of the canonical estrogen receptors alpha and beta (ERα and ERβ), and the G-protein coupled estrogen receptor (GPER, or GPR30).^1^ Canonical ERs are targeted directly to the nucleus, and interact with ER responsive elements (EREs) within the genome to regulate transcription, while GPER activation results in a cascade of posttranslational modifications in the cell that may also ultimately drive genomic responses.^2–4^

Estrogen receptor activation has been associated with changes in the abundance and activity of numerous enzymes involved in glucose and fatty acid energy metabolism, which result in significant sexual dimorphism in cardiac metabolic profiles. Of particular note, women exhibit greater cardiac fatty acid uptake and oxidation relative to men under both normal and pathophysiological conditions.^5^ Estrogen modulates the expression of cardiac metabolic proteins, and upregulates proteins that impact fatty acid metabolism, including PGC-1α and acyl-CoA dehydrogenases (ACADs).^6–8^

Consequently, the presence of estrogen has a significant impact on fuel substrate utilization in the heart.

The posttranslational acetylation of non-nuclear targets has emerged as a critical regulator of metabolic activity in the heart. In mitochondria, GCN5L1 and SIRT3 have been reported to increase and decrease, respectively, the acetylation status of enzymes that metabolize fatty acids and glucose.^9–15^ However, sex differences in the acetylation of metabolic proteins in cardiac mitochondria have not been investigated. We hypothesized that differences in the expression of GCN5L1 and SIRT3 between men and women may change the acetylation status and activity of enzymes involved in glucose and fatty acid metabolism, and that estrogen signaling may drive this process.

The studies presented here demonstrate that mitochondrial protein acetylation is increased in female mice relative to males, which is associated with sex-dependent elevations in GCN5L1 abundance. In addition, we show that estrogen directly increases GCN5L1 expression in human-derived cardiac cells, and that GCN5L1 is decreased in the hearts of postmenopausal women relative to younger women. The primary mechanism for estrogen-mediated GCN5L1 upregulation is identified as GPER activation, through a transcription-independent pathway. Finally, we determine that estrogen-dependent acetylation of MCAD is dependent on GCN5L1, and that loss of GCN5L1 results in diminished MCAD activity.

## METHODS

### Human tissues

Fresh human cardiac tissue samples were collected from the left ventricles of organ donors deemed not suitable for transplant, under a protocol approved by the University of Pittsburgh Committee for Oversight of Research and Clinical Training (CORID). Tissues were flash-frozen and stored at −80 °C until processing. Post menopause: range = 65-86 years, median = 69 years, N = 7. Pre menopause: range = 22-39 years, median = 36 years, N = 5.

### Animal care and use

All housing and experiments in mice were conducted in accordance with the guidelines established by the National Institutes of Health, and approved by the University of Pittsburgh Institutional Animal Care and Use Committee. Male and female C57BL/6J mice (aged 8-10 weeks) were purchased from The Jackson Laboratory, and maintained on a regular chow diet with a 12 h light/12 h dark light cycle.

### Cell culture and drug treatments

AC16 cells (a proliferating cell line derived from human cardiomyoctyes^16^) were purchased from Millipore. Stable GCN5L1 knockdown was generated as previously described.^17^ Cells were treated with 10 nM 17β-estradiol (E2), ICI 182, 780 (Fulvestrant; an ERα and ERβ antagonist with GPER agonist activity^18–20^), G-1 (a selective GPER agonist), G-36 (a selective GPER antagonist), MG-132 (a 26S protease inhibitor), and/or cycloheximide (CHX; a translation inhibitor).

### Mitochondrial Isolation

Mitochondrial fractions were purified from tissue and cells using the Qproteome Mitochondrial Isolation Kit (Qiagen) according to the manufacturer’s instructions. Briefly, samples were homogenized in cold Lysis Buffer, and centrifuged at 1000 *g.*

Supernatant containing the cytosolic fraction was discarded, and the pellet was re-suspended and processed in cold Disruption Buffer by shearing through a 25 g needle and syringe. Samples were centrifuged at 1000 *g*, the supernatant was collected, and centrifuged again at 6000 *g* to pellet the mitochondrial fraction. The pellet was washed in Storage Buffer, and then used for subsequent immunoblot or MCAD activity studies as described below.

### Immunoblotting

Tissue, cells, or purified mitochondria were lysed in 1% CHAPS buffer. Protein was quantitated using a BioDrop μLITE analyzer (BioDrop), and equal amounts were loaded on a 12% SDS-PAGE gel, before transfer to nitrocellulose membranes. Membranes were blocked using Odyssey blocking buffer and incubated in primary antibodies overnight (αTubulin, 1:1000, Cell Signaling; SIRT3, 1:1000, Cell Signaling; MCAD 1:1000, Cell Signaling; GCN5L1 1:500, generated as previously described^21^), followed by incubation at room temperature with fluorescent secondary antibodies for 1 h (800 nm anti-rabbit, LiCor). Bands were visualized using an Odyssey Imager, and quantitated using Image Studio Lite v 5.2 (LiCor).

### Quantitative RT-PCR

RNA was isolated from tissue or cells using RNEasy kit (Qiagen). RNA was quantified and 500 ng-1000 ng was used to generate cDNA using Maxima Reverse Transcriptase (Thermo Fisher). Quantitative PCR was performed using SYBR-Green (Thermofisher) and primers for GCN5L1 or SIRT3. GAPDH or PPIA were used for normalization.

### MCAD activity assays

MCAD activity was assessed using a DCPIP/PES-based assay as previously described. Briefly, DCPIP (50 μM), PES (2 mM), NEM (0.2 mM), KCN (0.4 mM), Triton X-100 (0.10%), and lysate were added to ice cold potassium phosphate buffer (0.1 M). MCAD substrate octanoyl-CoA was added to a final concentration of 40 μM, and then warmed to 37 °C for 5 min. Absorbance was read at 600 nm, and then normalized to protein concentration.

### Statistical Analysis

Statistical analyses were performed using GraphPad Prism 8.3. Student’s t-tests were used for simple comparisons between groups. One-way Analyses of Variance (ANOVA) was used to compare more than two groups, followed by post-hoc Student’s t-tests. For studies examining multiple time-points a two-way ANOVA was used with post-hoc Sidak’s multiple comparisons tests. A *P* value <0.05 was regarded as significant. All data are represented as the mean ± SEM.

## RESULTS

### Mitochondrial acetylation and GCN5L1 expression is increased in the hearts of female mice compared to male mice

To determine whether mitochondrial protein acetylation status is different between sexes, we isolated mitochondria from the hearts of male and female C57BL/6J mice, and immunoblotted for acetylated lysine residues. We observed a modest but significant increase in the intensity of bands in female mice compared to male mice (**Figure 1A**). We next examined whether the increase in mitochondrial acetylation is associated with changes in the expression of proteins that regulate mitochondrial protein acetylation. No changes were observed in the expression of the mitochondrial-targeted deacetylase SIRT3 (**Figure 1B**). However, a significant increase in both GCN5L1 mRNA and protein was observed (**Figures 1C and 1D**). Based on these data, we conclude that increased acetylation in female cardiac mitochondria is driven by increased GCN5L1 abundance.

**Figure 1:**
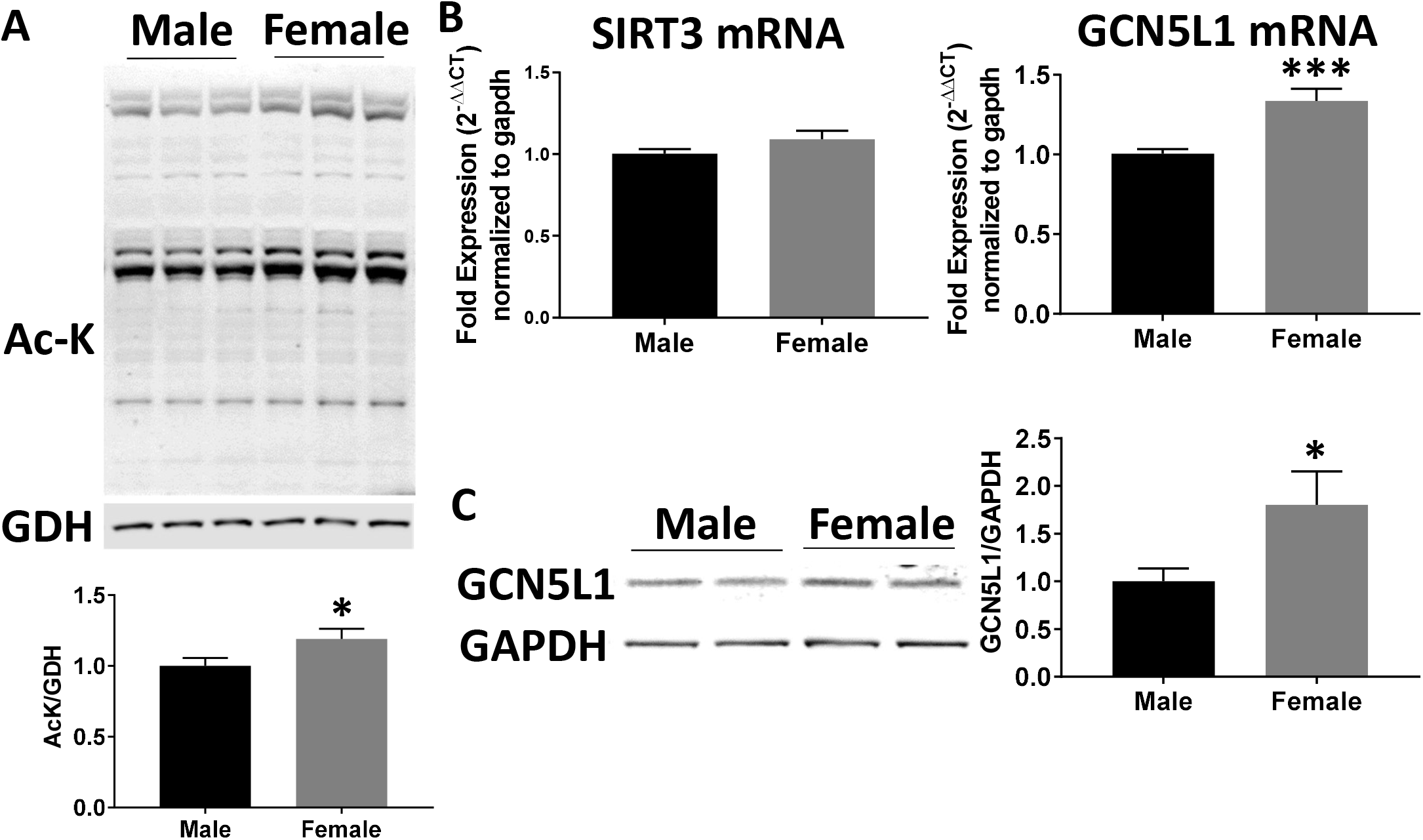
Mitochondrial protein acetylation and GCN5L1 are upregulated in female mice. A. Immunoblotting of mitochondrial lysate fractions from male and female mice demonstrate that females exhibit modestly but significantly higher levels of total protein acetylation. N = 10, B. GCN5L1, but not SIRT3, mRNA is significantly increased in female hearts relative to males. C. GCN5L1 protein levels are significantly increased in the myocardium of female mice compared to male mice. N = 10, * = p < 0.05, *** = p < 0.001 vs. male.

### Estrogen increases GCN5L1 expression in human cardiomyocytes via GPER

To determine if changes in GCN5L1 abundance are present in a clinically relevant setting, we analyzed heart tissue obtained from female patients of pre- and post-menopausal age. Immunoblotting revealed that cardiac tissues from women after menopause, when estrogen levels are lower, have a significantly lower GCN5L1 protein abundance (**Figure 2A**). As estrogen has been reported to mediate several of the sex differences observed in human myocardial tissue,^4^ we next determined whether estrogen induces GCN5L1 expression directly. Treatment of AC16 cells (derived from human ventricular cardiomyocytes)^16^ with 17-β estradiol (E2) resulted in significantly increased levels of GCN5L1 protein (**Figure 2B**). To determine whether signaling for increased GCN5L1 expression was through canonical estrogen receptors, we incubated AC16 cells with ICI 182,780 (Fulvestrant), a potent inhibitor of ERα and ERβ.

**Figure 2:**
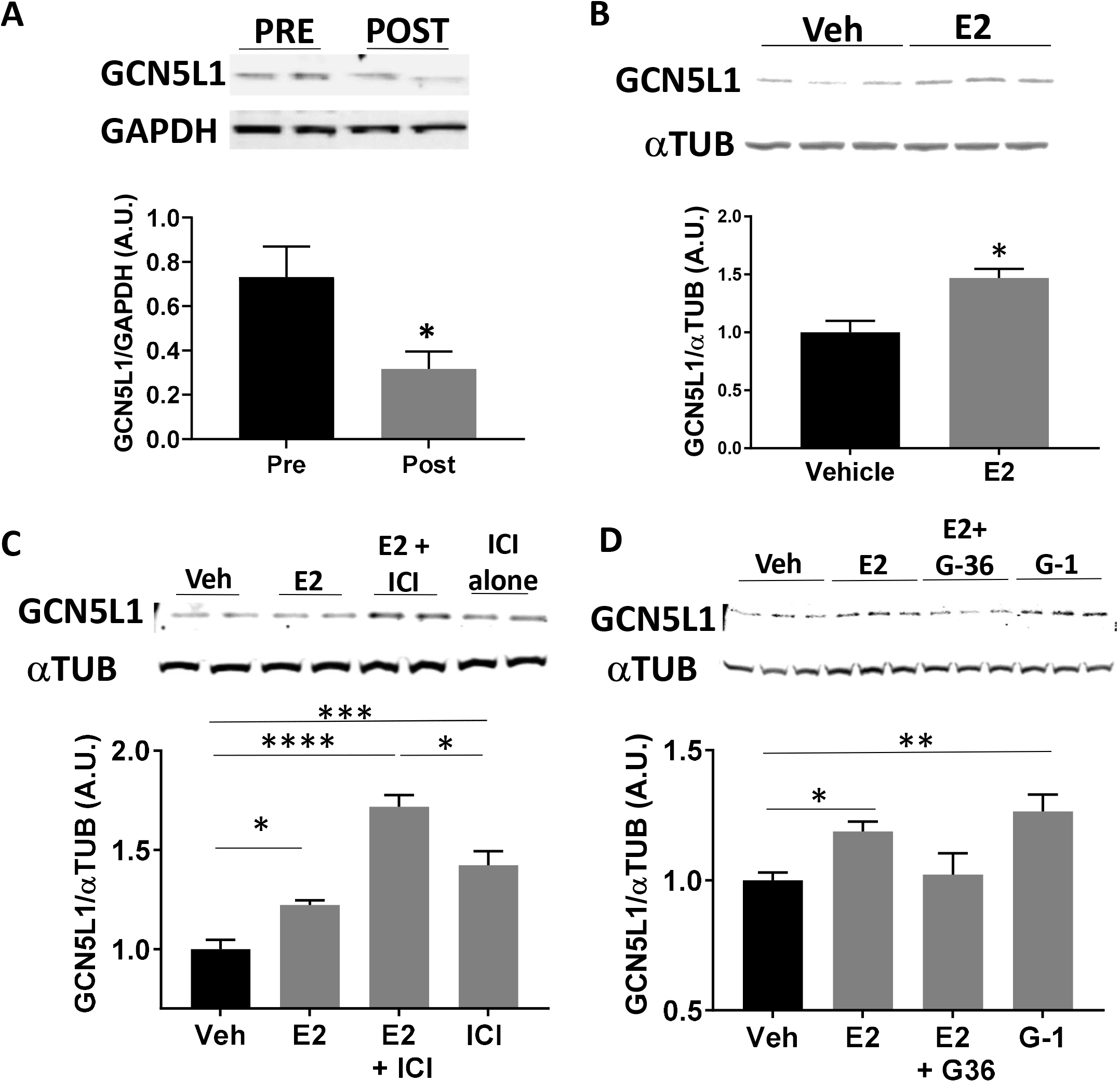
Estrogen drives GCN5L1 expression via GPER. A. Human tissues from female patients of pre-menopausal (PRE) or post-menopausal (POST) age. B. Human derived AC16 cells incubated with E2 show an increase in GCN5L1. C. GCN5L1 immunoblotting after incubation for 24 hours with E2 and or the ERa/ERb antagonist ICI 182, 780. N = 4, * = p < 0.05, **** = p < 0.0001 vs. vehicle. D. GCN5L1 is elevated after incubation for 24 hours with E2 and the GPER agonist G-1, and is blocked in the presence of GPER antagonist G-36. N = 5-6, * = p < 0.05, ** = p < 0.01 vs. vehicle.

Surprisingly, rather than blocking GCN5L1 induction, ICI 182,780 additively increased GCN5L1 levels, and produced a robust increase in GCN5L1 even in the absence of E2 (**Figure 2C**). ICI 182,780, in addition to blocking ERα and ERβ, also has been reported to act as a partial agonist for the G-protein coupled estrogen receptor (GPER).^19,22,23^ We therefore examined whether GPER played a role in estrogen-mediated GCN5L1 induction using the GPER agonist G-1, and the GPER antagonist G-36. Incubation with G-1 significantly increased GCN5L1 protein abundance, while G-36 blocked E2-mediated GCN5L1 expression (**Figure 2D**). From these data, we conclude that GCN5L1 abundance is increased by estrogen exposure via GPER-mediated signaling.

### Estrogen drives GCN5L1 translation

To understand the mechanism of GPER-induced GCN5L1 elevations, we monitored mRNA levels in AC16 cells after exposure to E2 or G-1 using qPCR. Surprisingly, no change in mRNA was observed at any of the time points measured, suggesting that E2 control of GCN5L1 expression occurs downstream of transcription (**Figure 3A**). To determine whether GCN5L1 protein elevation was due to an estrogen-induced reduction in protein degradation, the effects of 26S proteasomal inhibitor MG132 on GCN5L1 expression were evaluated. It was expected that if estrogen signaling elevates GCN5L1 levels by reducing protease activity, a blockade of protease activity would normalize protein levels in vehicle-treated cells, and the difference observed in E2-treated cells would disappear. Data showed that this was not the case, and rather MG132 amplified the increase in protein observed in the presence of E2 or G-1 (**Figure 3B**). We next examined whether estrogen alters GCN5L1 mRNA translation. To test this hypothesis, E2 and G-1 treated AC16 cells were incubated with the translational inhibitor cycloheximide (CHX). CHX treatment effectively normalized GCN5L1 levels, indicating that differences in protein expression may be attributed to GCN5L1 translational regulation (**Figure 3C**).

**Figure 3:**
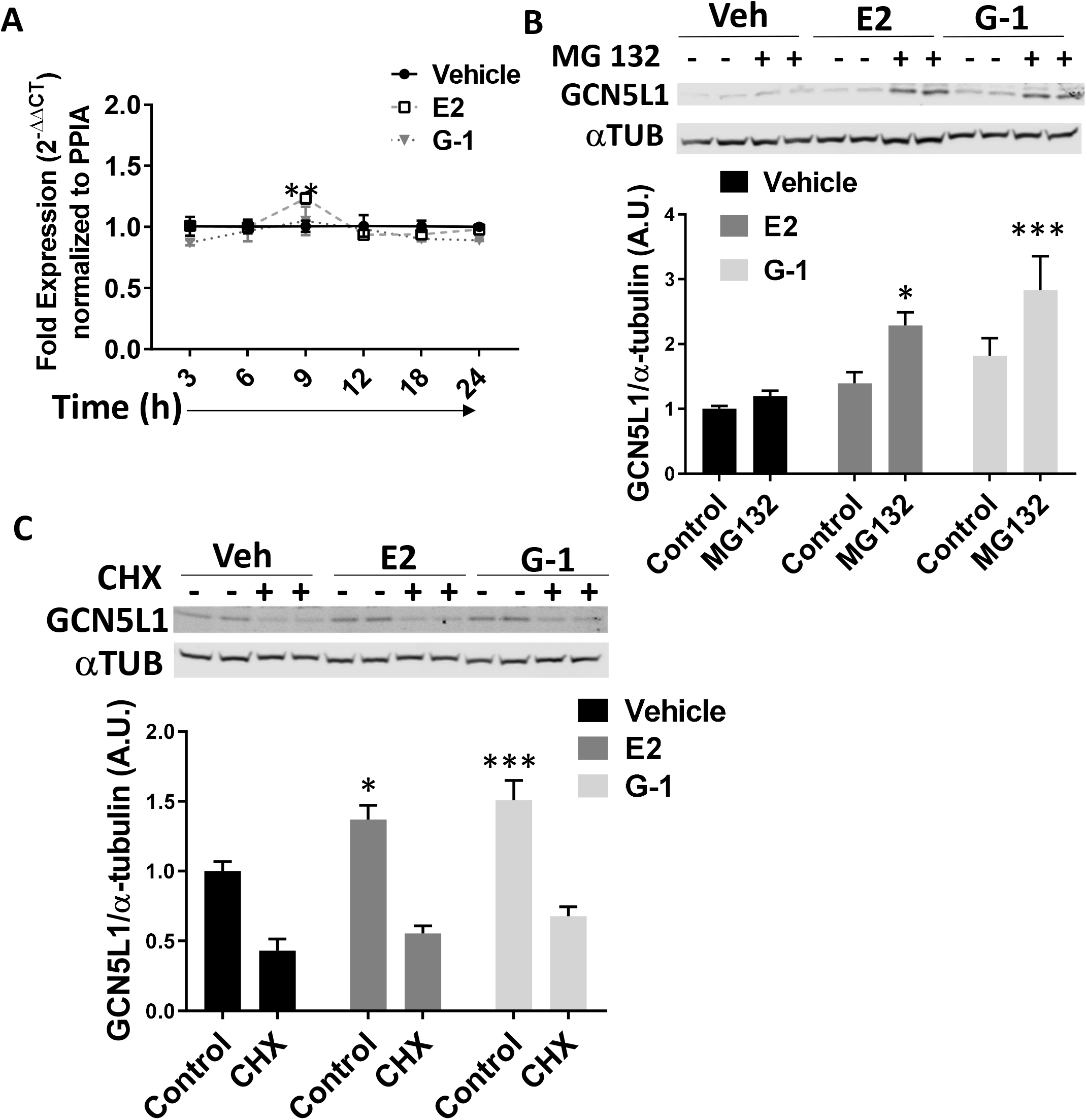
Estrogen promotes GCN5L1 translation. A. Expression of GCN5L1 mRNA levels determined by qPCR after treatment with E2 or G-1. N = 3, ** = p < 0.01 vs. vehicle. B. GCN5L1 levels are significantly elevated by blocking 26S degradation, suggesting that GPER agonism does not increase GCN5L1 via impaired proteolysis. N = 4, * = p < 0.05, *** = p < 0.001 vs. control. C. GCN5L1 induction is blocked by CHX, a translation inhibitor. N = 4 * = p < 0.05, *** = p < 0.001 vs. control.

### Estrogen-induced acetylation and activation of MCAD is reduced when GCN5L1 levels are depleted

Cardiac GCN5L1 has been reported to mediate the acetylation and activation of acyl-CoA dehydrogenases (ACADs), which mediate fatty acid breakdown.^12,24^ Among the ACADs, MCAD has been repeatedly identified as a target regulated by estrogens in the heart.^6–8^ We therefore tested whether GCN5L1 may link estrogen receptor agonism to increases in MCAD acetylation and activity. Estrogen treatment resulted in increased MCAD activity in AC16 cells, which was blocked in GCN5L1-depleted cells (Figure 4A). When GCN5L1 was silenced, MCAD acetylation was significantly reduced in E2 and G-1 treated cells, relative to control cells under the same conditions (Figure 4B). These data suggest that E2- and GPER agonist-induced MCAD activation occurs via GCN5L1.

**Figure 4:**
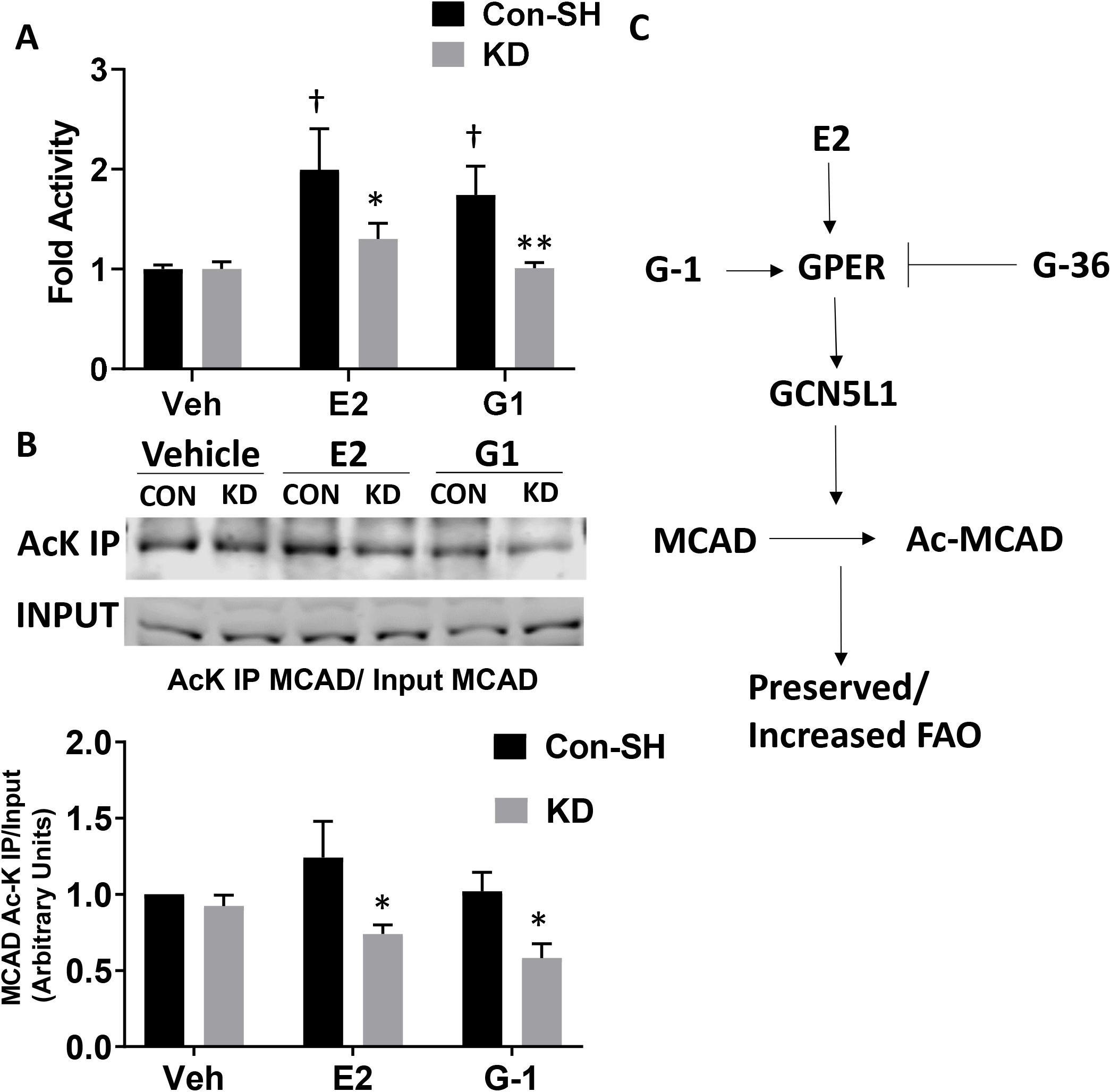
GCN5L1 is required for estrogen-mediated MCAD acetylation and activation. A. E2 and G-1 raise MCAD activity, while GCN5L1 shRNA knockdown blocks this effect. N = 8. B. Acetylation of MCAD is significantly reduced in the presence of E2 and G-1 when GCN5L1 is absent. N = 4 * = p < 0.05, ** = p <0.01 vs. control, † = p < 0.01 vs. vehicle. C. Schematic representing the hypothesized mechanism of action of estrogen on GCN5L1 in cardiomyocytes.

## DISCUSSION

Here we determined that mitochondrial protein acetylation, and GCN5L1 expression, are elevated in female mouse hearts compared to male. We find that estrogen upregulates GCN5L1 via GPER, and pharmacological block of GPER ablates induction. GCN5L1 is also elevated in pre-menopausal women, where estrogen levels are higher, compared to post-menopausal women. E2 and G-1 do not alter GCN5L1 gene transcription or proteasomal degradation; instead translational blockade prevents GCN5L1 induction. Finally, we determine that the loss of GCN5L1 blocks estrogen-mediated acetylation and activation of MCAD. These data point to a significant role for GCN5L1 in estrogen-mediated regulation of cardiac fuel metabolism (summarized in **Figure 4C**).

Significant differences in cardiac physiology and pathology between men and women are well established. Pre-menopausal women are largely protected from cardiovascular disease (CVD) compared to men, but this advantage is reduced with age.^25^ Estrogen loss has been suggested to be a major mediator of this effect. Postmenopausal women become more susceptible to left ventricular diastolic dysfunction, and hormone replacement therapy mitigates this effect.^26^ Ovarectomized mice and rats are similarly more susceptible to insults to the heart, including pressure overload,^27^ AngII induced hypertrophy,^28^ and diabetes-associated myofilament sensitization to calcium. ^29,30^ Ovariectomized mice exhibit a faster onset of obesity-driven heart failure, and show earlier signs of cardiac mitochondrial dysfunction, including elevated ROS production and swelling.^31^ Mice lacking estrogen receptors are found to be more sensitive to IR injury and hypertensive cardiomyopathy.^28,32^ Although less well-studied, testosterone also plays a role in driving sexual dimorphism between male and female individuals. Since estrogen was sufficient to reproduce an increase in GCN5L1 production, we have not evaluated the effects of testosterone. However, we cannot discount the possibility of an effect, and further studies are required to make this determination.

In recent years, GPER has taken a central role in our understanding of how estrogen impacts cardiac function and resiliency.^33^ GPER activation in cardiomyocytes lacking classical ERα and ERβ receptors was reported to alter intracellular calcium influx.^34^ In addition, GPER activation is associated with protection from ischemia-reperfusion injury, dependent on PI3K activation.^35^ Agonism of GPER with G-1 is reported to protect estrogen-deficient rats from LV remodeling.^36^ G-1 is also reported to inhibit the opening of the mitochondrial membrane permeability pore (mPTP)^37^, and reduces the upregulation of inflammatory cytokines TNF-alpha, IL-1beta, and IL-6.^38^ Our earlier observation that GCN5L1 protects the heart from I/R injury^17^ raises the possibility that GPER-mediated upregulation of GCN5L1 may be an additional mechanism by which estrogen protects the heart.

The acetylation status of several mitochondria-localized proteins has been reported to impact their function and stability. A key driver of acetylation status in mitochondria is the deacetylase SIRT3, which is expressed in myocardial tissue, and has been shown to regulate the function of several mitochondrial proteins involved in oxidative phosphorylation, membrane integrity, and redox homeostasis (See recent review by Chen et al.^39^). However, no difference in cardiac SIRT3 expression was identified between male and female mice, despite altered mitochondrial acetylation levels, suggesting that dimorphic effects in acetylation are due to GCN5L1 only.

GPER agonism drives translation and production of GCN5L1 protein, but at no time point after GPER activation was GCN5L1 message elevated. A recent publication by Lv et al. suggests that GCN5L1 abundance is regulated post-transcriptionally in diabetic kidney cells.^40^ We did not observe evidence that G-1 blocks the degradation of GCN5L1 by the 26S proteasome. Interestingly, there are reports of GCN5L1 acting as a translational coactivator to ERα in HeLa cells, binding directly to both the receptor and the corepressor element MTA1.^41^ That study did not evaluate a direct interaction between GCN5L1 and GPER, and although a mechanism by which GCN5L1 might interact directly with GPER was briefly considered, our immunoprecipitation studies did not support the direct binding of GPER to GCN5L1 (data not shown).

We demonstrate here that GCN5L1 is required for estrogen and GPER agonism to upregulate the acetylation and activity of medium-chain acyl-CoA dehydrogenase (MCAD) in cardiac-derived cells. MCAD is a mitochondria-localized enzyme that catalyzes the rate-limiting step in the β-oxidation of medium chain fatty acids, the α,β-dehydrogenation of fatty acyl-CoA. MCAD plays a key role in the progression of myocardial metabolic dysregulation induced by heart failure. TAC-mediated metabolic changes include the downregulation of MCAD,^42^ and gene delivery of MCAD to the heart protects against pressure overload-induced pathological remodeling.^43^ Subjecting mice to a high-fat diet was reported to increase contractile recovery after MI, concurrent with an increase in MCAD activity.^44^ Our laboratory and others have shown that high-fat feeding increases the activity of multiple ACADs through increased acetylation in response to GCN5L1 expression.^12^ In addition, MCAD has been reported previously to be regulated estrogen; E2 has been reported to upregulate or preserve cardiac MCAD expression via PGC-1.^6–8^ Studies of skeletal muscle biopsies report that women express higher levels of MCAD,^45^ and MCAD expression was reported to increase in men after treatment with E2.^46^ The studies reported here show for the first time that estrogen may also impact through posttranslational acetylation mediated by GCN5L1 expression.

In summary, we demonstrate here for the first time that GCN5L1 is upregulated by estrogen signaling in both mouse and human myocardium, and in cultured cardiomyocytes. The mechanism of upregulation is identified as GPER-mediated signaling. MCAD is identified as a target of acetylation and activation in the presence of E2 or GPER agonists, which is dependent on GCN5L1 expression. These findings shed new light on the role that posttranslational acetylation, mediated by GCN5L1, may play in the differences observed between men and women in cardiac metabolism.

## ACKNOWLEDGEMENTS

This work was supported by NIH/NHLBI R01 (HL132917 & HL147861) research grants to IS, and a UPMC CMRF Research Grant to JRM.

